# Proteome-wide inference of protein kinase regulatory circuits

**DOI:** 10.1101/703157

**Authors:** Brandon M. Invergo, Borgthor Petursson, David Bradley, Girolamo Giudice, Evangelia Petsalaki, Pedro Beltrao

## Abstract

Complex networks of regulatory relationships between protein kinases comprise a major component of intracellular signaling. Although many kinase-kinase regulatory relationships have been described in detail, these are biased towards well-studied kinases while the majority of possible relationships remains unexplored. Here, we implement data-driven, unbiased methods to predict human kinase-kinase regulatory relationships and whether they have activating or inhibiting effects. We incorporate high-throughput data, kinase specificity profiles, and structural information to produce our predictions. The results successfully recapitulate previously annotated regulatory relationships and can reconstruct known signaling pathways from the ground up. The full network of predictions is relatively sparse, with the vast majority of relationships assigned low probabilities. However, it nevertheless suggests denser modes of inter-kinase regulation than normally considered in intracellular signaling research.

## Introduction

Cells continually respond and adapt to environmental stimuli. They employ sophisticated protein networks to propagate, amplify and subsequently quench these signals efficiently. A common mechanism of relaying information from one protein to another is through reversible post-translational modifications (PTMs). Protein phosphorylation by kinases is one of the principal and best-studied PTMs. It plays a major role in cellular processes such as growth, division and differentiation (Acosta-Jaquez et al., 2009; Basson, 2012; Rhind and Russell, 2012).

Many protein kinases are themselves regulated by phosphorylation, giving rise to complex networks of kinase-kinase regulatory relationships. An accumulation of biochemical knowledge has produced consensus maps of several protein-kinase signaling pathways, which have been deposited in databases such as Reactome (Fabregat et al., 2018), KEGG (Kanehisa et al., 2016) and SIGNOR (Perfetto et al., 2015). Kinase-kinase and other kinase-substrate relationships have also been annotated in databases such as PhosphoSitePlus and Phospho.ELM (Dinkel et al., 2011; Hornbeck et al., 2015). However, bias in the study of kinase regulatory relationships has left the majority of the kinase-kinase interaction space largely unexplored (Invergo and Beltrao, 2018). Similar biases have been reported for protein-protein interaction databases (Gillis, Ballouz, and Pavlidis, 2014). Subsequent, unbiased analyses have found protein interactions to be ultimately more evenly spread across the proteome than previously indicated (Rolland et al., 2014), and the same is likely to be true for kinase signaling.

These biases can have serious impacts on systems-level analyses of signaling pathways. There is, therefore, a clear need for new, unbiased methods for finding kinase-kinase regulatory relationships. Existing methods for data-driven reconstruction of signaling networks are generally designed for data that has been produced for the study of a specific pathway (e.g. via perturbation experiments) and typically benefit from the incorporation of prior knowledge about that pathway into the model (see, e.g., Hill et al. (2016) and Invergo and Beltrao (2018)). Given the study bias inherent to this approach, these methods are less likely to provide insight into broader patterns of protein kinase regulation, especially of understudied kinases or cross-module signaling. However, recent advances in high-throughput phosphoproteomics, through liquid-chromatography tandem mass-spectrometry (LC-MS/MS) and other technologies, show promise in the inference and analysis of signaling networks (Babur et al., 2018; Rudolph et al., 2016; Terfve et al., 2015).

Alternatively, computational methods can be used to prioritize future experiments in an unbiased manner. Here, we propose a machine-learning approach to estimate the probability of a regulatory relationship between two kinases, as well as to predict the sign (inhibiting or activating) of the regulation. We produced the predictions by combining phosphoproteomic and transcriptomic data with kinase substrate-sequence specificity models and a recently produced predictor of phosphosite functional impact (Ochoa et al., 2019). Together, these data allow us to make inferences even for kinases that lack any substrate annotations. The resulting network of predicted kinase-kinase regulatory relationships is highly modular and partitions into several clusters that reflect known functional associations, while suggesting denser modes of inter-regulation and feedback than typically assumed.

## Results

### Regulatory relationships can be identified by similar phosphorylation patterns at functional phosphosites and kinase coexpression

Since many protein kinases are regulated by phosphorylation, we first measured correlations in phosphorylation at regulatory phosphosites between kinases. If regulatory sites on two kinases show similar patterns of phosphorylation, one of the kinases might be responsible for regulating the other’s activity. We assessed correlations of phosphosite quantifications in two large-scale phosphoproteomic experiments (Mertins et al., 2016; Wilkes et al., 2015). Regulatory functionality has only been assigned to a small subset of kinase phosphorylation sites, so to improve coverage, we employed a recently produced computational predictor of phosphosite functionality (Ochoa et al., 2019). This provided us with a score from 0.0 to 1.0 for each site, with higher values indicating a stronger prediction of functional impact of phosphorylation (“functional sites”).

We found that kinase-kinase regulatory pairs often exhibit co-phosphorylation patterns at functional phosphosites. For example, mitogen-activated protein kinase 3 (MAPK3) is known to regulate the activity of ribosomal protein S6 kinases (Mérienne et al., 2000; Smith et al., 1999; Zhao, Bjorbaek, and Moller, 1996). Indeed, we found strong correlation between functional sites T202 on MAP kinase 3 and T577 on S6K-alpha-3 (RPS6KA3); meanwhile, no such correlation was found for atypical MAP kinase 4 (MAPK4), which has no known regulatory relationship with S6 kinases (Figure 1a). We quantified this relationship for each pair of sites between two kinases by producing a phosphosite “coregulation score”, in which the log-transformed *p*-value of the correlation is scaled by the two sites’ functional scores (Figure 1a). In both phosphoproteomic experiments, kinase-kinase regulatory pairs tend to exhibit higher maximum coregulation scores than pairs with no previously annotated relationship (Figure 1b).

**Figure 1:**
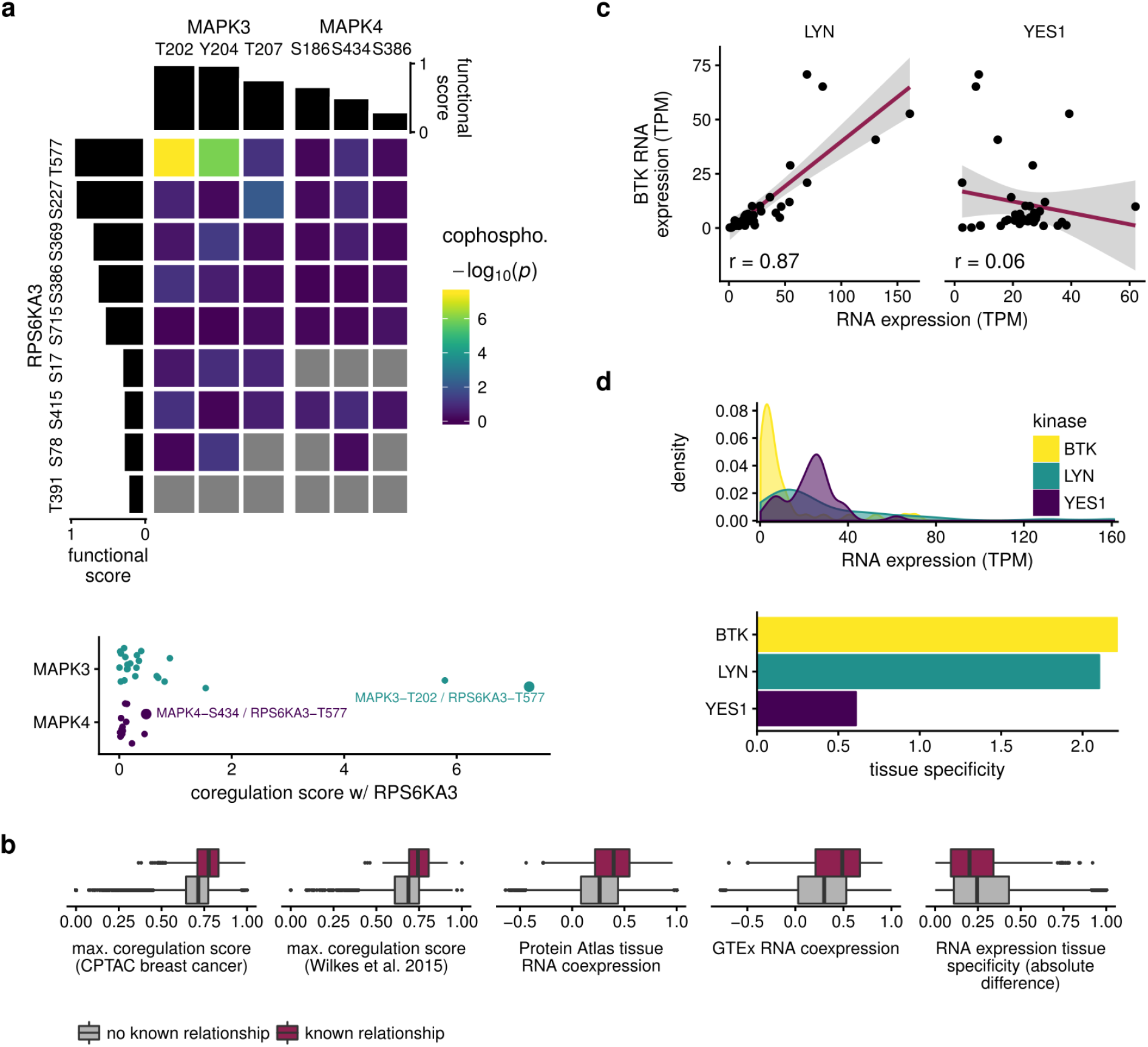
Correlations in phosphorylation at regulatory sites or in tissue expression patterns are predictive of kinase-kinase regulatory relationships. a) Top: Kinase MAPK3 exhibits significant cophosphorylation patterns at functional sites with RPS6KA3, a known substrate. The same patterns are not observed for MAPK4. Gray cells indicate missing values. Bottom: Combining cophosphorylation significance and site functional scores provides an estimator of coregulation. b) Phospho-coregulation, tissue coexpression and tissue specificity can discriminate known cases of kinase-kinase regulation from unannotated cases. c) The RNA transcripts encoding SRC-family kinase LYN and known substrate BTK show similar patterns of expression, while the expression of SRC-family kinase YES1, not known to regulate BTK, is unrelated. d) Top: Kernel-density estimates of the distributions of expression values across tissue samples for *BTK*, *LYN* and *YES1*. Bottom: Tissue-specificity of RNA expression was quantified as the skewness of the kernel-density distributions. Here, *YES1* is more broadly expressed than the tissue-specific *LYN* and *BTK*.

We next used two RNA-seq data sets (GTEx Consortium, 2013; Uhlén et al., 2015) to test whether kinase coexpression is indicative of regulatory relationships. For example, if we consider the regulation of tyrosine-protein kinase BTK by Src-family protein kinases, we see a clear positive correlation between *BTK* expression and that of *LYN* (encoding tyrosine-protein kinase Lyn, a known regulator) (Cheng, Ye, and Baltimore, 1994; Park et al., 1996; Rawlings et al., 1996). No such correlation exists for *YES1* (tyrosine-protein kinase Yes, which is not known to regulate BTK) (Figure 1c). In general, we found higher coexpression between pairs of kinases where a regulatory relationship exists than for those without any annotated relationship in both expression data sets (Figure 1b).

We also found that tissue specificity, as represented by the skewness of expression values across tissue samples, is further indicative of kinase regulatory relationships. Continuing from the previous example, we can see that *BTK* and *LYN* both have skewed expression profiles (high expression in a few tissues), whereas *YES1* has relatively even expression across tissues (Figure 1d). If we consider the absolute difference between tissue specificities for pairs of protein kinases, we find that pairs with regulatory relationships tend to have more similar expression profiles than those with no annotated relationship (Figure 1b).

### Linking sequence specificity to phosphosite functional impact identifies direct regulation of protein kinase activity

Kinases show preferences for phosphorylating some substrates over others, determined by the specific phosphoacceptor residue and a surrounding amino-acid sequence. By characterizing this specificity in a position-specific scoring matrix (PSSM), we can score a kinase’s potential for directly phosphorylating a putative substrate. However, we also wanted to determine, in an unbiased way, whether high-scoring substrate sites also tend to have regulatory effects. To achieve this, we employed the discounted cumulative gain (DCG) metric often used in the evaluation of information retrieval systems (Järvelin and Kekäläinen, 2002), wherein we treated a PSSM as a phosphosite “search function” and the functional score as a phosphosite “relevance metric.”

Only 140 protein kinases had sufficient numbers of known substrate sites to build confident PSSMs. We have recently shown that proteins within the same kinase family tend to show similar specificity, which can be attributed to conserved specificity-determining residues (SDRs) within their protein-kinase domains (Bradley, Vieitez, et al., 2018; Bradley and Beltrao, 2019). We thus investigated this as a means to assign PSSMs to kinases with insufficient substrate annotations. We first estimated the minimum residue similarity necessary across 10 kinase SDRs to make accurate PSSM assignments. We found that a SDR similarity of at least 0.8 (based on the BLOSUM62 amino-acid substitution matrix) is needed to make assignments that are significantly better than a random assignment (Figure 2a). Nevertheless, this method of assignment did not substantially improve upon simply assigning a family-wise, composite PSSM (Figure 2b). Based on these results, we increased the coverage of kinases with PSSMs by assigning to under-annotated kinases a family-wise PSSM where available (*n* = 208) or otherwise one via SDR similarity (*n* = 14), bringing the total number of protein kinases with specificity profiles to 362 (Figure 2c).

**Figure 2:**
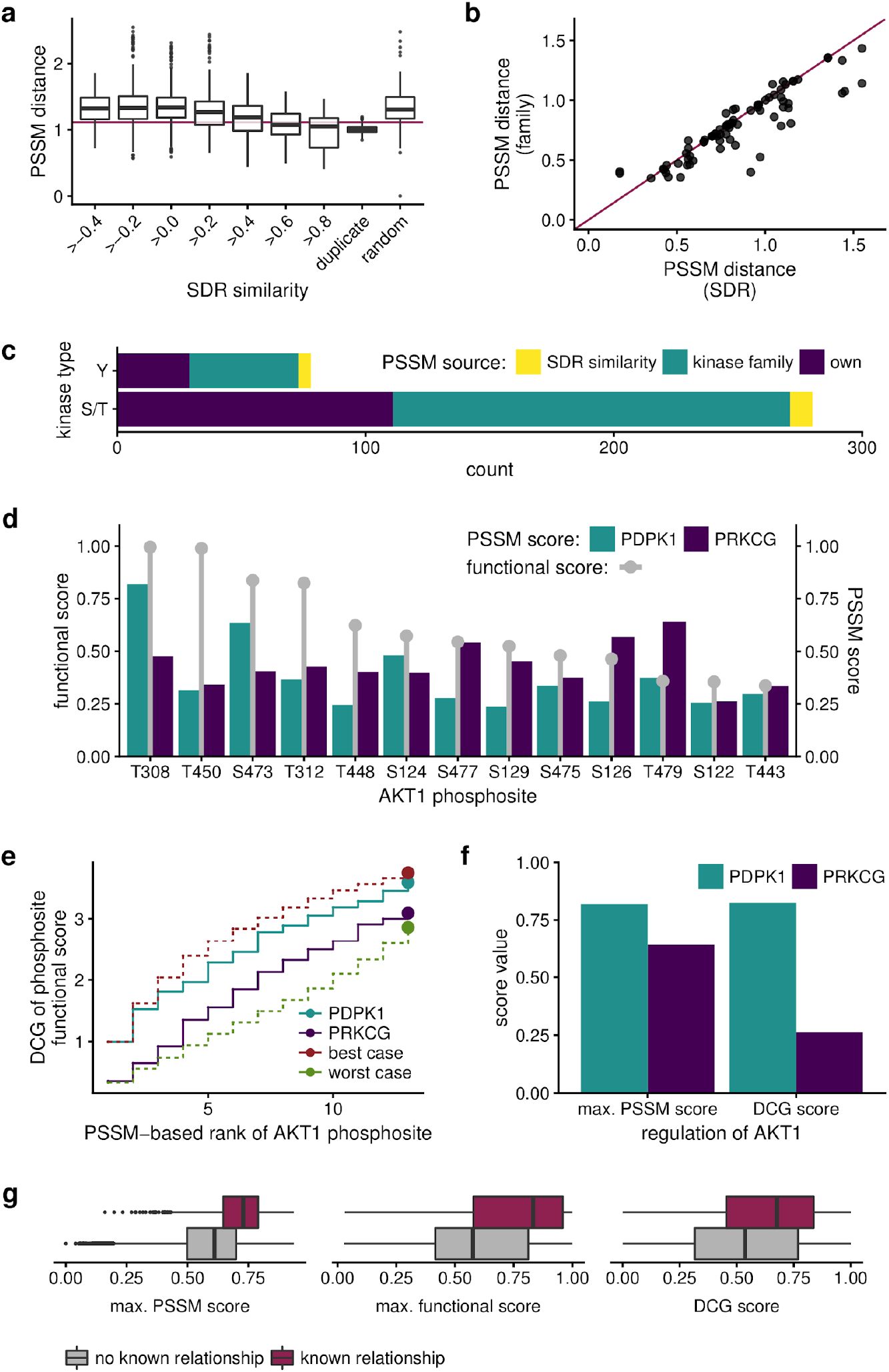
Kinase-kinase regulatory relationships can be predicted from sequence specificity and phosphosite functional scores. a) Similar kinase specificity-determining residues also indicate similar PSSMs. The red line indicates the 97.5th percentile of the distribution of distances between cross-validation PSSMs using different subsets of a kinase’s annotated substrates. At an SDR similarity of at least 0.8, over 50% of assigned PSSMs are less than this distance from their true values. b) Assigning family-wise, composite PSSMs to unannotated kinases achieves similar, if not better, performance than SDR-based assignment. c) Numbers of PSSMs by source (own annotations = 140, by family = 208, by SDR similarity = 14). d) PSSMs locate functional sites on substrates with differing performance. Here, the PSSM of PDPK1, a known regulator of AKT1, scores highly for sites with high functional scores, while that of PRKCG does not. e) Discounted cumulative gain (DCG) quantifies the potential for a kinase to phosphorylate a putative substrate at its functional sites. f) Although both PDPK1 and PRKCG have similar maximum PSSM scores for phosphorylating AKT1, only PDPK1 achieves a high DCG. g) Maximum PSSM score, maximum substrate-site functional score and DCG all discriminate known regulatory relationships from unannotated ones.

Linking PSSM predictions to phosphosite functional scores via the DCG is best illustrated by an example. RAC-alpha serine/threonine-protein kinase (AKT1) has several phosphosites, a few of which have high functional scores. We consider two potential regulators: 3-phosphoinositide-dependent protein kinase 1 (PDPK1), a known regulator; and protein kinase C gamma type (PRKGC), not known to regulate AKT1. Some of AKT1’s sites with the highest functional scores also score highly with PDPK1’s PSSM, whereas PRKCG’s PSSM favors sites with low functional scores (Figure 2d). These relationships can be quantified and visualized via the DCG: substrate sites are ranked by PSSM score and a cumulative sum of their functional scores is calculated, wherein each successive site contributes a smaller fraction of its functional score (Figure 2e). We can see that, although the two protein kinases achieve similar maximum PSSM scores, only PDPK1 produces a high DCG (Figure 2f).

As would be expected, we found that the PSSMs of known regulators tend to score highly for at least one of their substrate’s phosphosites (Figure 2g, left panel). Furthermore, simply having a substrate site with a high functional score, indicating that the substrate is amenable to regulation by phosphorylation, can be predictive of a regulatory relationship (Figure 2g, center panel). Linking these two metrics across all sites on the substrate via the DCG, we produced a score that could discriminate true regulatory relationships (Figure 2g, right panel).

### Protein sequence and structure discriminate phosphosites that induce or inhibit kinase activity

Phosphorylation events can lead to different regulatory outcomes for the substrate kinase, potentially inducing or inhibiting its enzymatic activity. Knowing these regulatory effects is essential to understanding the flow of information across complex networks of regulatory relationships. Thus, we sought to infer the “signs” (activating or inhibiting) of regulatory relationships from data.

To do so we first evaluated how phosphorylation at a specific site is likely to affect a given kinase’s activity. We found particular discrimination for sites within phosphorylation hotspots of the protein-kinase domain (Strumillo et al., 2019): sites within hotspots tend overwhelmingly to be activating (i.e. within the kinase activation loop) (Figure 3a, 1st panel). Interestingly, when considering the sites’ positions within the domain, we found that most inhibitory sites are N-terminal (Figure 3a, 2nd panel), whereas they tended to be more C-terminal in the overall protein (Figure 3a, 3rd panel). Lastly, we also observed that activating sites tend to be in more structured regions of the protein and inhibitory sites are more likely to be disordered, although 50% of all inhibitory sites still were predicted to be in structured regions (Figure 3a, 4th panel).

**Figure 3:**
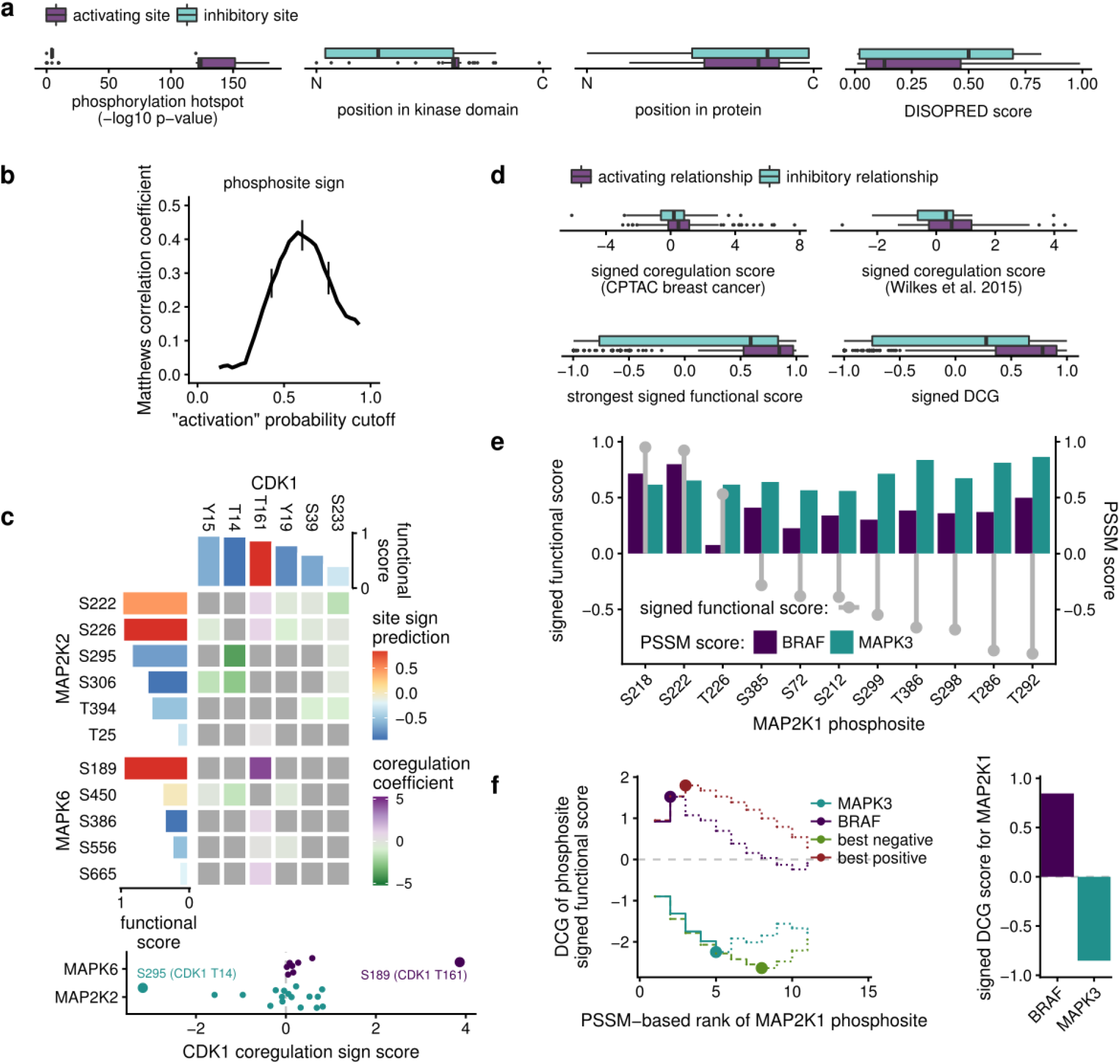
Evidence of regulatory sign (activating vs. inhibiting) can be uncovered in a data-driven manner. a) The regulatory sign of a single phosphosite can be discriminated by using structural information: whether the site is in a phosphorylation hotspot; where the site is within the protein-kinase domain (N = N-terminal, C = C-terminal); the relative position of the site within the protein (N = N-terminal, C = C-terminal); and whether the site is in a disordered region;. b) Matthews correlation coefficients for different posterior-probability cutoffs for the predictor of phosphosite regulatory sign. The cutoff (above which a site or relationship would be declared to be “activating”) that maximizes the coefficient discriminates best between inhibitory and activating sites or relationships. Error bars represent 95% confidence intervals. c) Modifying the phospho-coregulation score to account for predicted phosphosite sign and correlation sign can produce protein-level predictions of regulatory sign. Here, CDK1 is shown to have an activating relationship with MAPK6 and an inhibitory relationship with MAP2K2. Gray cells indicate missing or removed values. d) The signed variants of the coregulation score, functional score, and DCG all discriminate between inhibitory and activating kinase-kinase regulatory relationships. e) Accounting for predicted phosphosite sign can assess the propensity of a kinase to phosphorylate activating or inhibiting sites: BRAF’s PSSM scores highly for activating sites on MAP2K1, while MAPK3 scores highly for inhibitory sites. f) A modified DCG for signed functional scores correctly assigns BRAF as an activator of MAP2K1 and MAPK3 as an inhibitor. Because there are more inhibitory sites on MAP2K1, a full DCG would be negative in most cases (dotted lines). Instead, we take the most extreme value visited by the sum (solid lines).

We then trained a predictor of phosphosite regulatory sign using these features (Table S1) via the Bayesian Additive Regression Trees (BART) method. Cross-validation of the model showed consistently good performance, with a maximum mean Matthew’s correlation coefficient of 0.42 at a cutoff of 0.58 (posterior probabilities lower than the cutoff are declared to reflect inhibitory functionality), indicating overall good sign-classification performance (Figure 3b). Adjusting these posterior probabilities by the highest-performing cutoff provided us with a sign score for all phosphosites in our data set, with negative scores indicating a prediction of inhibition of activity and positive scores predicting activation (Table S2).

### Kinase regulatory sign can be inferred from phosphosite sign and interaction evidence

With phosphosite sign predictions in hand, we aimed to predict the signs of kinase-kinase regulatory interactions. Returning to the coregulation of functional phosphosites, we tested the consistency of the observed phosphoproteomic correlation with the sign of the phosphorylating kinase’s regulatory site. If phosphorylation of an inhibitory site on a kinase is anticorrelated with that of an activating site on a putative substrate, then the evidence would suggest that the kinase positively regulates the substrate’s activity. On the other hand, no direct-regulation scenario would explain a positive correlation between these sites.

For example, cyclin-dependent kinase 1 (CDK1) shows strong evidence of negative coregulation with dual specificity mitogen-activated protein kinase kinase 2 (MAP2K2), reflecting its role in inhibiting MAP kinase kinases (Rossomando et al., 1994) (Figure 3c). CDK1 also activates Mitogen-activated protein kinase 6 (*MAPK6*) (Tanguay, Rodier, and Meloche, 2010) and, indeed, we find a strong positive correlation between two activating sites on these kinases (Figure 3c). Overall, we found that the signed coregulation score was able to discriminate between activating and inhibitory kinase regulatory relationships in both phosphoproteomic datasets (Figure 3d, first and second panels).

We also adapted our DCG methodology after applying our sign predictions to the site functional scores. Thus, we now asked whether a kinase’s PSSM tends to find relevant activating sites or inhibitory sites. For example, dual specificity mitogen-activated protein kinase kinase 1 (MAP2K1) is activated by serine/threonine-protein kinase B-raf (BRAF) (Alessi et al., 1994; Macdonald et al., 1993; Papin et al., 1995) and is inhibited in negative feedback by its downstream substrate, mitogen-activated protein kinase 3 (MAPK3) (Eblen et al., 2004). We found that, indeed, B-raf has specificity towards MAP2K1’s activating sites while MAPK3 is specific towards the inhibitory sites (Figure 3e). We then calculated a DCG on the signed functional scores, taking the most extreme value visited by the sum (Figure 3f). This method provides a positive value for BRAF and a negative value for MAPK3, as expected. Overall, both the signed functional score and the signed DCG score could discriminate well between activating and inhibitory relationships. However, predictions for inhibitory relationships overall were less reliable (Figure 3d, third and fourth panels).

### Predicting a global network of kinase regulatory relationships

We combined the above evidence into two predictors via machine learning. The edge predictor predicts whether a kinase-kinase regulatory relationship exists. The sign predictor predicts whether a given relationship induces or inhibits the substrate’s kinase activity.

For training and validating the edge predictor, we retrieved from the OmniPath metadatabase (Türei, Korcsmáros, and Saez-Rodriguez, 2016), a list of annotated relationships with at least two source databases supporting them, comprising 825 interactions in all. Because it is more difficult to prove the absence of a regulatory relationship, there is a lack of annotations for genuinely false relationships. We assumed that, in the space of all possible kinase-kinase interactions, regulatory relationships are rare. Therefore, a randomly selected pair of kinases is unlikely to show any regulatory relationship. We thus assessed the features described above for their predictive power on a validation set consisting of the annotated positive cases and random “negative” subsets of the remaining space of putative interactions.

Overall, each of the edge-predictor features (Table S3) exhibited limited but measurable predictive power. We visualized this by the receiver operating characteristic (ROC) curve, comparing true-positive and false-positive rates as the score-cutoff for declaring a regulatory relationship is lowered; and by similarly assessing precision and recall across cutoffs (Figure 4a). Maximum PSSM score performed the best, with a mean area under the ROC curve (AUC) of 0.742 (*σ* = 0.007, *n* = 100) (Figure S1a). The remaining features had mean AUC values of less than 0.7. We also noted that the precision decayed rapidly with lower cutoffs.

**Figure 4:**
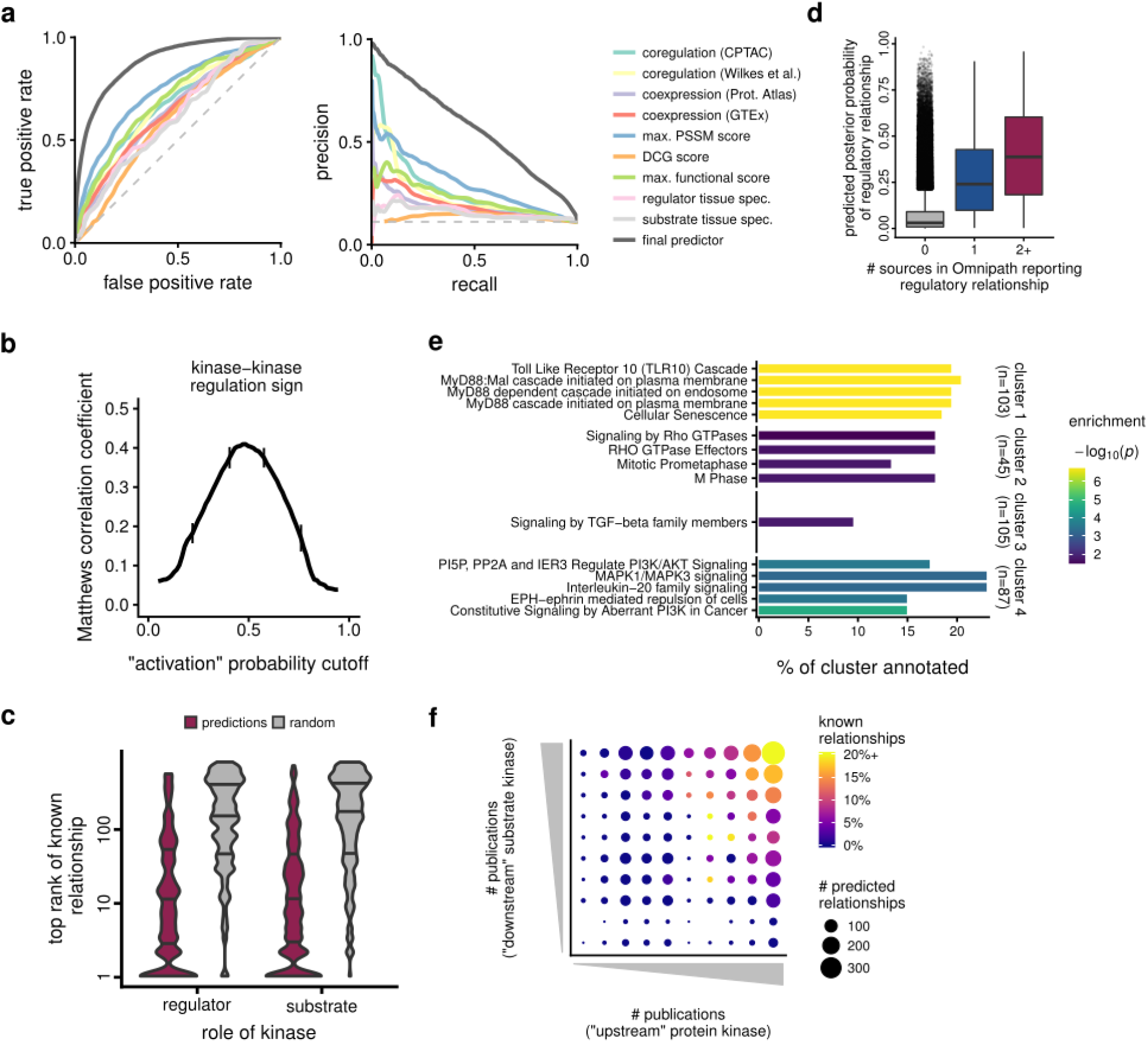
Combining data-driven predictors of kinase-kinase regulatory relationships. a) ROC and precision-recall curves of each feature and the final edge predictor. See also Figure S1a. b) Matthews correlation coefficients for different posterior probability cutoffs for the sign predictor. Error bars represent 95% confidence intervals. c) Annotated regulatory relationships for each kinase tend to rank highly among the predictions, when considering the kinase as either a regulator or a substrate. Lines indicate quartiles. 50% of kinases had a known regulatory relationship in the top ten predictions, which is significantly better than random expectation. See also Figure S2b. d) Previously annotated relationships supported by only one source in OmniPath score similarly to those supported by two or more sources (used in our training set), further validating our predictions. e) Clusters identified on the regulatory sub-network at a posterior probability cutoff of 0.5 are significantly enriched in annotations for unique sets of pathways. See also Figures S1b–c. f) The predicted network expands upon the annotated network, especially for understudied protein kinases. See also Figure S2a.

We then combined these features into the edge predictor using the BART method (Chipman, George, and McCulloch, 2010) (Table S4). We first performed 3-fold cross-validation on the model 20 times using different random iterations of the training set (Figure 4a). The resulting models had a mean AUC of 0.884 (*σ* = 0.009, *n* = 60), representing a significant improvement over the individual features (Figure S1a). The robustness of the cross-validation results also assured that the model was not over-fitting the training set.

We applied the same BART method to the regulatory sign features (Table S5) produce the sign predictor (Table S4). We trained the model using regulatory signs annotated in OmniPath and evaluated it through cross-validation. Overall, performance was similar to the underlying site-level predictor described above, with a mean maximum Matthews correlation coefficient of 0.42, however confidence intervals over the cross-validation were narrower for kinase-level predictions than they were for site-level predictions (Figure 4b). The maximum correlation occurred at a cutoff of 0.484 (i.e., the probability above which we would declare regulation to activate the substrate).

We next considered whether known, annotated relationships tend to rank highly among our edge predictions for each kinase. We found that 50% of kinases had a known regulatory relationship within the top 10 of our predictions (Figure 4c). The top ranks were significantly better than expected, based on random, per-kinase permutations of the scores (one-sided Wilcoxon rank sum test, regulator: *W* = 5818.5, *p* < 1 × 10^−6^; substrate: *W* = 7385.5, *p* < 1 × 10^−6^).

To further evaluate our model, we looked at how well it predicted interactions that were not included in the positive set due to being supported by only one source in OmniPath (*n* = 293). This provided a completely new, external validation set. These interactions had significantly higher prediction scores than unannotated regulatory interactions, however they were generally lower than the high-confidence set (one-sided Wilcoxon rank sum test vs. unannotated: *w* = 6 × 10^7^, *p* < 1 × 10^−6^, vs. high-confidence set: *W* = 87073, *p* < 1 × 10^−6^; Figure 4d).

We also noted several high-probability predictions that, while not being annotated in OmniPath, have direct or plausible support in the literature. For example, we predict receptor tyrosine-protein kinase erbB-2 (ERBB2/HER2) to activate ephrin type-A receptor 2 (EPHA2) (edge probability 0.94, sign probability 0.79). These two oncogenic kinases form a complex and in a mouse model of breast cancer they appear to cooperate to promote tumor progression (Brantley-Sieders et al., 2008), however no direct regulatory relationship has yet been described. We also predict that the closely related tyrosine protein kinases Fer (FER) and Fes/Fps (FES) activate HGF receptor (MET) with equal probabilities (edge probabilities 0.92, sign probabilities 0.80). In fact, activation of MET by FER has previously been reported (Fan et al., 2016), however this relationship is not annotated in OmniPath and thus was not present in our training and validation set. As a final example, we predict tyrosine protein kinase ABL1 to activate focal adhesion kinase 1 (PTK2/FAK1) (edge probability 0.92, sign probability 0.81). While, to our knowledge, no such regulatory relationship has been described previously, FAK1 plays an important role in acute lymphoblastic leukemia characterized by constitutively active ABL1, and its phosphorylation was speculated to be “likely augmented by the direct action of activated ABL1 itself” (Churchman et al., 2016).

We next assessed the topology of regulatory relationships using a sub-network of high-confidence predictions (probability greater than 0.5), consisting of 340 kinases and 4339 regulatory relationships (representing less than 2% of all possible relationships). We first applied a cluster-detection algorithm to an undirected variant of this network (retaining the higher-probability relationship when two kinases were predicted to regulate each other, producing 3716 undirected edges). Four clusters consisting of more than 10 kinases each were detected (103, 45, 105 and 87 kinases, respectively; Table S6). This division of the network had a modularity of 0.325, which was significantly higher than expected given the modularity of randomized networks with the same degree distribution (*µ* = 0.197, *σ* = 0.00339; *p* < 0.001; Figure S1b). To determine if these clusters reflected known biological associations, we tested each one for enrichment in pathway annotations from Reactome (Fabregat et al., 2018). Each cluster was enriched in annotations for at least one distinct Reactome pathway, indicating that the network successfully identified clusters of physiologically related kinases (Figure 4e; Table S7). We also assessed how related the pathways associated with each cluster were, using the average number of reactions between the proteins of two pathways as a proxy for relatedness. We found that the pathways associated with the same cluster were more closely related to each other than to those associated with other clusters (*p* < 1 × 10^−6^, *W* = 5.8 × 10^5^, Wilcoxon rank sum test; Supplemental Figure S1c).

Because we set out to overcome the study bias inherent to signaling pathway annotations, we checked for relationships between kinase connectivity on the high-confidence network and kinase publication counts (Figure 4f; kinase publication counts were retrieved from Invergo and Beltrao (2018)). Interactions between kinases in the top three publication-count deciles (more than 95 publications) accounted for only 31% of the network. Conversely, 589 regulatory relationships were predicted between pairs of kinases in the bottom 50% of publication counts (fewer than 40 publications each).

Overall, only 7% of the relationships in the high-confidence network are annotated in databases. Although the number of previously annotated interactions is dwarfed by novel predictions, a significant proportion of this can be accounted for by the relative sparsity of annotated relationships for under-studied kinases. Restricting the network to highly-studied kinases largely resolves this (Figure S2a). However, this can also be explained in part by the persistence of study bias in our results, as can be seen in a significant correlation between publication count and top prediction-rank of known relationships (Figure S2b; Spearman’s rank correlation, as regulator: *ρ* = *−*0.34, *p* < 1 × 10^−6^; as substrate: *ρ* = *−*0.29, *p* < 1 × 10^−6^).

### Reconstructing signaling pathways from kinase regulatory predictions

We next investigated whether our data-driven, signed kinase-kinase regulatory predictions were able to reconstruct known pathways. We applied an edge probability cutoff of 0.5 and a sign cutoff of 0.5. We started by choosing well-studied kinases that are functionally related to AKT1 (Figure 5a). Between these kinases, we successfully recovered all but one annotated relationship, the regulation of ribosomal protein S6 kinase beta-1 (RPS6KB1). Six predicted relationships are not present in database annotations. Sign predictions generally fail on a per-substrate basis. For example we predict all regulations of RPS6KB1 to be inhibitory, while those that have been annotated are activating. Our predictions perform even better when considering MAPK signaling, again recovering all but one previously annotated edge, but with only one erroneous prediction of an annotated sign (Figure 5b).

**Figure 5:**
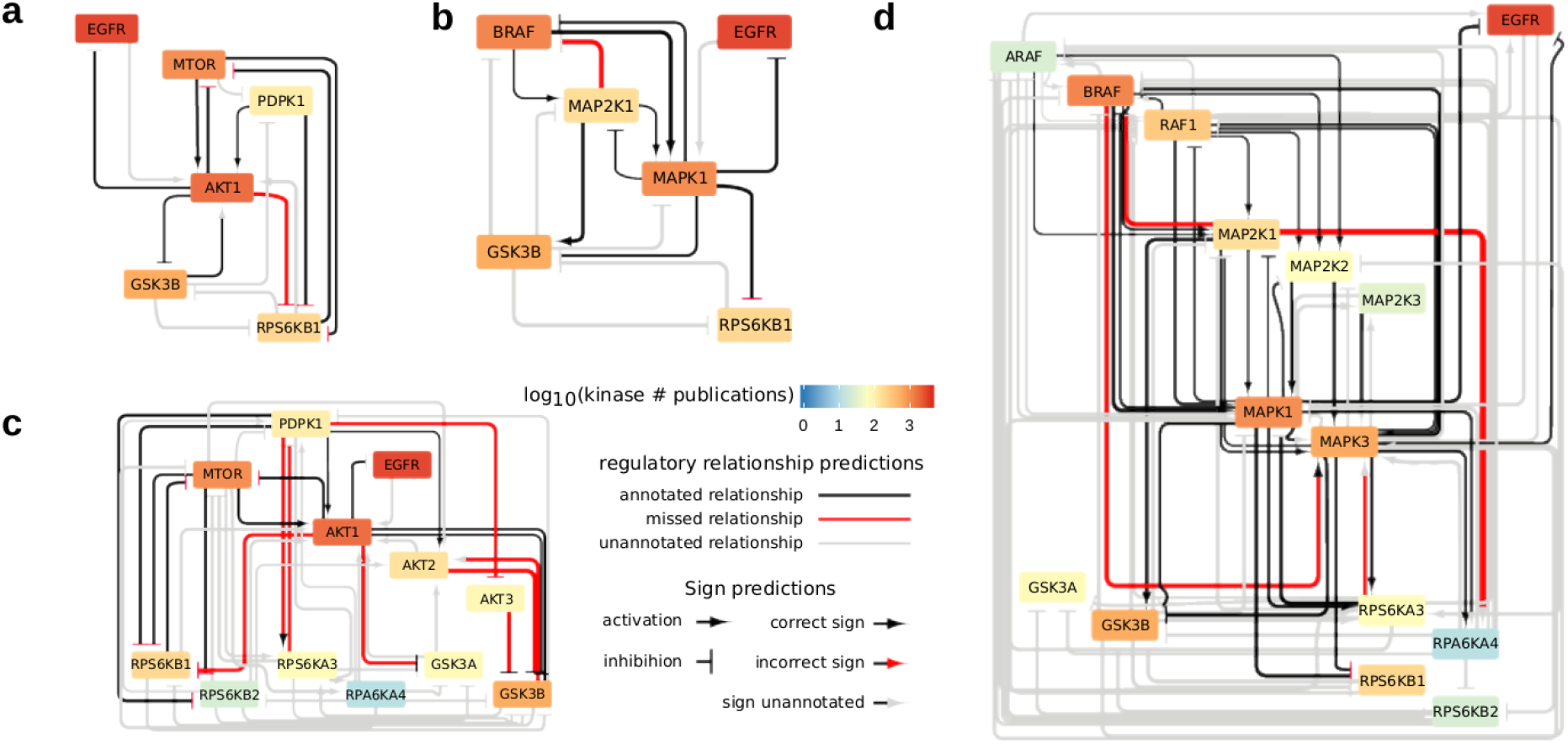
Our data-driven predictor reconstructs known signaling pathways “from scratch”. a) A reconstruction of AKT1 signaling at a probability cutoff of 0.5 and “activating” sign cutoff of 0.5. Black edges are correctly recovered. Red edges failed to be predicted. Gray edges are unvalidated predictions. Arrowheads indicate the predicted regulatory sign: arrows indicate activation and bars indicate inhibition. Black arrows are correctly predicted, red arrows are incorrectly predicted and gray arrows are unvalidated. Node colors indicate the number of publications associated with the kinase. b) Similar performance is seen in reconstructing MAP kinase signaling from our predictions. c–d) Including more kinases, particularly understudied kinases, greatly increases the number of unvalidated edges while also predicting complex modes of regulatory feedback.

If we begin to include other paralogs of these kinases, which tend to be less well-studied, we quickly accumulate predictions for previously undescribed relationships. For example, we predict many modes of inter-regulation between S6 kinases and glycogen synthase kinases. On the other hand, we fail to predict several regulatory relationships involving RAC protein kinases AKT2 and AKT3 (Figure 5c). Expanding the MAPK signaling network is more successful again, with the core signaling events being recovered between RAFs, MAP2Ks and MAPKs, including correct sign prediction, while also filling in the network of interactions for the less well-studied A-Raf (ARAF) and MAP2K3 (Figure 5d). Both of these examples demonstrate that the predicted networks quickly become difficult to assess when more than a few kinases are included, particularly those with fewer annotations. However, extrapolating from the overall performance and the success on smaller networks, our results suggest that this complexity is inherent to kinase signalling networks.

## Discussion

The task of experimentally testing all possible kinase-kinase relationships in order to produce an unbiased network is daunting. We have thus taken a data-driven approach to predict these regulatory relationships. We do not suggest that these predictions can replace established methods for confirming regulatory relationships. However, they can be used to reduce the vast space of possible relationships under consideration in order to form credible hypotheses and to prioritize experiments, particularly for under-studied kinases.

Previous efforts to produce kinome-scale inferences have depended on scaffolding data-driven predictions to existing protein networks. For example, Rudolph et al. (2016) derived signaling pathways through a network diffusion technique with phosphoproteomic data on a literature-derived protein-protein interaction network. However, such analyses are strongly impacted by the biases present in the existing networks (Gillis, Ballouz, and Pavlidis, 2014; Rolland et al., 2014). To our knowledge, there has only been one other attempt to predict kinase regulatory sign (Hernandez et al., 2010). The authors inferred signs from quantitative phosphoproteomic data on a literature-derived kinase network, in which the method in part depended upon the connectivity of the kinases on this network. However, missing or erroneous annotated relationships could have major impacts on the results. By building our network from the ground up with data, we largely removed these biases from our predictions and were nevertheless successful in reconstructing known, signed regulatory networks. We only retain study bias from using annotated substrates in the construction of our kinase-specificity models. This could be resolved with high-throughput methods to measure kinase specificity profiles (see, e.g. Imamura et al. (2014)).

Many factors can affect the nature of a kinase-kinase regulatory relationship and each such relationship will be unique, owing to the particular properties of the kinases involved. Thus, making generalized predictions about them is inherently difficult. Nevertheless, some features are fundamental, such as regulation by phosphorylation. To this end, the performance of the predictions via identifying patterns of phosphorylation will improve with more data. Given the importance of PSSMs in our results, there is a clear need for producing robust PSSMs for every kinase in order to prune indirect regulatory effects. As for correlative methods on phosphoproteomic data, many conditions are needed to confidently discriminate the phosphoregulation of over 500 kinases. Importantly, large-scale phosphoproteomics experiments are needed across a more diverse array of tissues or cell lines to properly capture the activities of more tissue-specific kinases. Because we only used data from experiments using the breast cancer cell line MCF7, many kinases were not represented in the phosphoproteomic data. Furthermore, the use of data derived from cancer cell lines might introduce errors in the resulting network since cancer initiation and progression disrupt intracellular signaling (Deribe, Pawson, and Dikic, 2010).

We assumed in the construction of our predictor that the true network is sparse, and indeed we assign to 75% of all possible relationships posterior probabilities of less than 0.09, far below any probability cutoff that we considered. Nevertheless, even at stringent cutoffs, isolating a subnetwork of more than a few kinases produces a denser topology of regulatory relationships than is typically considered for kinase signaling. It is possible that this is an artifact of not considering cellular context (e.g. protein expression or cellular localization). There is also an unavoidable accumulation of false-positives as more predictions are considered. Despite these caveats, our results suggest that the kinase regulatory network is richer in feedback and cross-module regulation than expected based on the current, biased view of kinase pathways. Further developments in experimental approaches for unbiased kinase regulatory network reconstruction are needed to confirm the predicted modularity and density of regulatory relationships in kinase signalling networks.

## Methods

### Data

We defined the human kinome as the list of 504 human proteins identified as protein kinases in the UniProt/Swiss-Prot Protein Knowledgebase, *pkinfam* (accessed 8 November 2017 at https://www.uniprot.org/docs/pkinfam Quantitative phosphoproteomic data was retrieved from two publications. The first included phosphosite quantifications of 213 phosphosites for 100 kinases across 22 kinase-inhibitory conditions in MCF7 cells (Wilkes et al., 2015). The second quantified 1537 phosphosites on 193 kinases across 83 breast tumor samples (Mertins et al., 2016). Tissue RNA expression data for protein kinases were retrieved from the GTEx project (GTEx Consortium, 2013) as provided by Expression Atlas (E-MTAB-5214, timestamp 26 April 2018) (Papatheodorou et al., 2018). We furthermore retrieved tissue RNA expression data from the Human Protein Atlas project (accessed from https://www.proteinatlas.org/ 1 December 2017) (Uhlén et al., 2015). Lists of human phosphosites, kinase substrates and kinase regulatory sites were retrieved from the Phospho-SitePlus database (accessed 1 May 2018) (Hornbeck et al., 2015). Amino acid frequencies in the human proteome were derived from the UniProt proteome database (UniProt Consortium, 2018).

### Protein kinase specificity models

#### Constructing kinase specificity models

We estimated kinase specificity through the construction of position-specific scoring matrices (PSSMs) from the amino acid sequences around known substrate sites (+/−7 residues), omitting autophosphorylation sites. We required at least 10 known substrates in order to build a PSSM for a given kinase, resulting in PSSMs for 140 protein kinases. In order to reduce the influence of redundant sequences on the construction of the matrices, we employed a position-based sequence-weighting method (S. Henikoff and J.G. Henikoff, 1994).

Given a set of *n* ≥ 10 substrate amino-acid sequences, *S* = {*S*_1_, *S*_2_, …, *S*_*i*_, …, *S*_*n*−1_, *S*_*n*_}, where *S*_*i*_ = {*S*_*i*1_, *S*_*i*2_, …, *S*_*i*14_, *S*_*i*15_} and *S*_*ij*_ represents the amino acid at position *j* of sequence *i*, we give a weight to each of amino acid *a* at position *j* as follows:

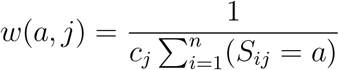

where *c*_*j*_ is the number of unique amino acids found in position *j* among the substrates in *S*. Next, a weight is calculated for each sequence as the sum of its position-specific residue weights:

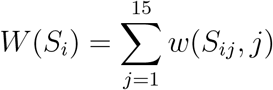

Finally, each sequence weight was normalized by the sum of all sequence weights:

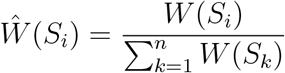

A 20 × 15 PSSM can then be constructed as follows. First, we construct matrix *r*, such that entry *r*_*aj*_ contains the weighted count of amino acid *a* at position *j* across the sequences in *S*:

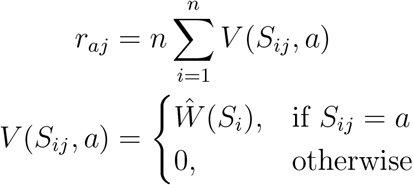

There is a non-zero probability of observing each residue at a position in the sequence, however at small sample sizes, we are unlikely to accurately estimate low-probability occurrences. To overcome this, we added pseudocounts based on proteome-wide amino-acid frequencies in a position-specific manner (Henikoff and Henikoff, 1996). For each column *j* in the PSSM, we select a number of pseudocounts, *B*_*j*_, to add:

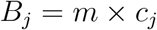

where *m* is a tune-able parameter and *c*_*j*_ is defined as above. Thus, empirically constrained positions (e.g. the +1 position for proline-directed kinases) will receive fewer pseudocounts, and thus lower baseline probabilities of observing other residues, than highly variable positions. We found that our results were not strongly dependent on *m*, so we fixed it at 1. A 20 × 15 matrix of pseudocounts, *b*, was then calculated as follows:

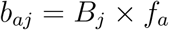

where *f*_*a*_ is the occurrence frequency of amino acid *a* in the proteome. This allows us to derive an empirical matrix of probabilities, *p*, of observing amino acid *m* at position *j*:

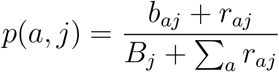

The final PSSM was arrived at by calculating the *log*_2_ fold-change of *p_aj_* versus the proteomewide amino acid frequencies:

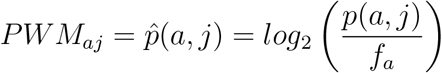

### Assigning PSSMs to protein kinases

In order to increase our coverage of specificity profiles to include protein kinases with few or no known substrates, we assigned to them either composite, family-wise PSSMs or PSSMs of protein kinases with similar specificity determining residues (SDRs) (Bradley, Vieitez, et al., 2018). For each protein kinase family, we constructed a family-wise PSSM as described above using known substrates of all kinases in the family, as defined by the KinBase resource (http://kinase.com/web/current/kinbase/). This family-wise PSSM was then assigned to any member of the family for which we could not construct a unique PSSM. PSSMs were assigned to 209 protein kinases in this manner.

Finally, for the remaining protein kinases for which no family-wise PSSM was available, we attempted to assign a PSSM based on SDR similarity. Towards this end, 10 kinase domain positions were selected, representing residues known to covary with kinase specificity and that are proximal (<4Å distance) to the kinase substrate at the active site (Bradley, Vieitez, et al., 2018). For a given pair of kinases, sequence similarity across the 10 SDRs was calculated by summing BLOSUM62 substitution scores for each position. An ‘SDR similarity’ score was then calculated by dividing this sum by the maximum possible score across the 10 SDRs, such that identical kinases would yield a similarity score of 1.0.

As represented in Figure 2a, the relationship between SDR similarity and PSSM distance was explored systematically to decide upon an SDR similarity threshold to use for PSSM assignment. For this purpose, SDR similarity scores and PSSM distances were calculated for all possible pairwise comparisons of kinases with known specificity. Here, similarity between PSSMs was quantified using the Frobenius distance, which represents the sum of squared element-wise distances between matrix values, followed by taking the square-root (Ellis and Kobe, 2011). For reference, pairwise Frobenius distances were also calculated for PSSMs of the same kinase by subsampling known target sites of a given kinase, using a sample size of 25 targets sites (corresponding to the median number of target sites used for PSSM construction). The distribution of all possible pairwise distances among these ‘duplicate’ PSSMs had a median of 1.00 and a 97.5th percentile of 1.10 (Figure 2a, red line). We interpret PSSM distances below the 97.5th percentile to represent kinases with the same active site specificity. An SDR similarity threshold of 0.8 was therefore selected as more than half of kinase pairs above this value have PSSM distances below the 1.10 threshold. For PSSM assignment, targets from the most similar kinase(s) in the human kinome were selected, provided the SDR similarity score was above 0.8. We assigned PSSMs to a further 14 kinases through this method. For all PSSM comparisons, the phospho-acceptor column (P0: S/T/Y) was not used when calculating the Frobenius distance.

The predictive performance of family-based and SDR-based PSSM predictions was compared in Figure 2b. For every kinase of known specificity, a PSSM was assigned using the family-based and SDR-based approaches described above, and then the Frobenius distance between empirical and predicted PSSMs was calculated for both prediction methods.

### Scoring phosphosites with PSSMs

For each directed protein kinase-kinase relationship, we scored each known phosphosite on the substrate kinase using the upstream kinase’s PSSM. For the +/−7 motif sequence around a given phosphosite (omitting the phosphosite itself), we calculated the PSSM score, *s*, as:

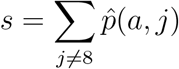

In order to make scores comparable between kinases, we then calculated a normalized score, *s*, against the minimum and maximum scores attainable with the PSSM:

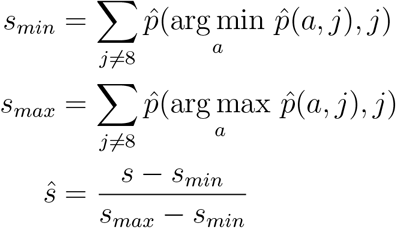

### Phosphosite functional scores

Predictions of functional relevance of phosphosites were retrieved from Ochoa et al. (2019). The predictions were made on a variety of phosphosite structural, evolutionary and biochemical features. As the predictions were originally made on a strictly defined set of phosphosites derived from a reanalysis of a set of high-throughput phosphoproteomics experiments, not all of the phosphosites available in the PhosphoSitePlus database were represented. We *log*_10_ -transformed the raw scores and normalized them against the minimum and maximum values to arrive at functional scores valued between 0.0 and 1.0, with larger scores reflecting a greater expectation of a functional impact of phosphorylation at that site.

### Linking PSSMs to phosphosite functional scores via Discounted Cumulative Gain

We assessed a kinase’s potential to phosphorylate a putative substrate at sites of likely functional relevance by linking the kinase’s PSSM to the substrate’s phosphosite functional scores via a Discounted Cumulative Gain calculation (DCG). In effect, we treat the PSSM as a “search function” and we employ the functional scores as relevance scores to determine how well a PSSM “finds” functional sites. For each substrate phosphosite with a functional score available, we calculate the PSSM score *s* as above. Next, the *n* sites are ranked by *s* in descending order, producing an associated ordering of functional scores *F* = {*F*_1_, *F*_2_, …, *F*_*i*_, …, *F*_*n*−1_, *F*_*n*_}. The DCG for this kinase-substrate pair is then calculated as:

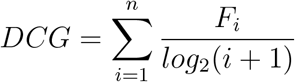

Sites with higher PSSM scores, and thus lower rank *i*, contribute larger fractions of their functional scores to the sum. The DCG will be highest, then, if sites with high functional scores tend to have high PSSM scores.

In order to make DCG scores comparable between different kinase-substrate pairs, we normalized each score by the minimum and maximum possible DCG scores for the substrate. The minimum DCG for a substrate can be found by sorting the sites in ascending order of their functional scores; likewise, the maximum can be found by sorting the sites in descending order of their functional scores. Thus, the normalized DCG is:

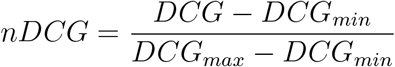

### Protein kinase coexpression and expression specificity

Coexpression of protein kinases across tissues in the GTEx and Protein Atlas RNA expression datasets was calculated via Spearman’s correlation after setting missing values to 0.0.

The tissue specificity of each kinase was calculated by assessing the skewness of its distribution of Protein Atlas expression values (in transcripts per million, or “TPM”) across the samples, defined as

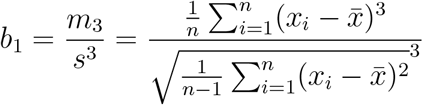

where *x* is the kinase’s set of expression values across all tissues, *x* is the sample mean expression value, *s* is the sample standard deviation, and *m*_3_ is the third central moment of the distribution. Skewness was calculated using the “e1071” package for R (https://cran.r-project.org/package=e1071).

### Correlation between phosphorylation levels across different samples and conditions

We assessed coregulation of a pair of protein kinases by measuring the correlation between phosphorylation of their regulatory phosphosites across conditions (Wilkes et al., 2015) or tissue samples (Mertins et al., 2016) of phosphoproteomic experiments. Both experiments consisted of a table of *log*_2_ fold-changes for each quantified phosphosite across the conditions or samples, measuring the relative intensities of the phosphosite as detected by mass spectrometry under each condition or sample versus a reference. The data for each experiment was quantile normalized by condition or sample (Bolstad et al., 2003).

Within an experiment, for each pair of phosphosites on two protein kinases, we calculated the correlation between fold-changes of the sites across all conditions or samples for which a quantification was available, where at least five such conditions or samples existed. We first removed any conditions in which one of the kinases was under chemical inhibition. The correlation was calculated using Spearman’s rho and a *p*-value estimated for the correlation via the asymptotic *t* approximation. *p*-values were then −*log*_10_ -transformed; in the case that the estimated *p* was 0, we set the final value to 6. This value was then scaled by the functional scores of both sites such that only two sites with high functional scores and a high phosphorylation correlation would have a high final coregulation score. We then took the maximum such score across all site pairs as the final coregulation score for the kinase pair. Finally, the coregulation scores for all kinase pairs were normalized according to the maximum and minimum values of all pairs.

### Prediction of regulatory relationships

#### Training and validation set

We retrieved a high-confidence set of known, directed kinase-kinase regulatory relationships from OmniPath (Türei, Korcsmáros, and Saez-Rodriguez, 2016) (fetched Jan 22, 2018 via the Python API). To ensure the quality of the relationships we only used those that were supported by at least two sources, providing a “positive” set of 825 relationships. It is more challenging to define a “negative” set of regulatory relationships, given the difficulty in unequivocally demonstrating a lack of regulation under all conditions. However, we assume that regulatory relationships are rare and that, given a random pair of kinases, there is unlikely to be a regulatory relationship between them. Working under this assumption, we constructed negative sets by randomly sampling from the space of possible relationships. To further reflect the presumed sparsity of the true network, we chose to construct a negative set that was 8 times larger than the positive set, which provided a slight improvement in prediction performance. This value was arbitrarily chosen to balance the diminishing performance boost from increasing negative set size with the rapidly increasing memory resources required to perform the training computations.

#### Feature validation

We evaluated the performance of the following features for predicting protein kinase regulatory relationships: maximum PSSM score, maximum substrate phosphosite functional score, DCG, phosphoproteomic coregulation scores, tissue RNA coexpression, and regulator and substrate tissue expression specificity. Each feature was evaluated 100 times against a randomly sampled two-thirds of the positive set and an 8-fold larger randomized negative set.

#### Model Training and Prediction

In order to build a final predictive model of kinase-kinase regulatory relationships, we employed the Bayesian Additive Regression Trees (BART) method (Chipman, George, and McCulloch, 2010). Briefly, BART is a “sum of trees” method, in which a series decision trees are fit to the data and used to classify data. Each tree consists of binary decision nodes reflecting a decision based on one of the features, e.g. “max. PSSM score > 0.75” or “GTEx coexpression < 0.3”. The terminal nodes of the tree contain values which, once selected, contribute to the final classification value; in a sum-of-trees model, the decision values from each tree are summed to produce a value upon which this final classification is made. The BART method, in particular, uses a fixed number of trees, on which it places regularizing priors that ensure that each tree is a “weak learner”, i.e. each tree contributes a small fraction of the final classification value. It does this by restricting the tree depth, shrinking terminal leaf nodes to the median, and adding noise to avoid over-fitting. Trees are fit to the data through Bayesian approaches to estimating the parameters, such as Markov-Chain Monte Carlo (MCMC) back-fitting (Chipman, George, and McCulloch, 2010).

We applied the BART model to our data as implemented in the R package “bartMachine” version 1.2.3 (Kapelner and Bleich, 2016). A notable extension of the original method provided by bartMachine is to incorporate data missingness into predictions (Kapelner and Bleich, 2015). For example, a missing phosphoproteomic coregulation value might be informative as it would indicate that phosphosites on the two kinases were never detected in the same conditions by the mass spectrometer, and thus a decision tree node asking “is the coregulation score missing?” can contribute to the final classification. We used this feature by enabling the “use_missing_data” and “use_missing_data_dummies_as_covars” parameters and disabling the “replace_missing_data_with_x_j_bar” and “impute_missingness_with_x_j_bar_for_lm” parameters. These settings in effect disable any imputation of missing data and produce new “dummy” variables that indicate whether a value is missing, which can then be incorporated in the decision trees. Model hyperparameters including the number of trees were determined using the built-in 5-fold cross-validation routine provided by the “bartMachineCV” function.

In addition to the quantitative features listed in the previous section, we also included the kinase types (serine/threonine versus tyrosine) for the regulating kinase and the substrate kinase as additional features. We evaluated the BART model on these features using the full “positive” training set and random “negative” training sets as outlined above. To this end, we performed 20 iterations of 3-fold cross-validation, using a different random “negative” set each iteration. We evaluated the true-positive rate, false-positive rate, the precision (positive predictive value) and the recall (sensitivity) of the model based on the calculated posterior probabilities assigned to the validation set. Performance metrics were calculated using the R package ROCR (Sing et al., 2005).

In order to produce our final classifications, we trained 100 different BART models to the training set, each with a different random instantiation of the “negative” set. Each model was then used to produce a posterior probability of a regulatory relationship for all kinase-kinase pairs. Finally, we took the mean of the 100 posterior probabilities for each relationship as the final classification score.

### Network clustering and pathway enrichment

The resulting network was divided into clusters using the method of Blondel et al. (2008) as implemented by the R package “igraph” in the function “cluster_louvain” (Csárdi and Nepusz, 2006). This is a heuristic method that identifies clusters by optimizing modularity. The algorithm can be divided into two steps: first, a cluster is assigned to each node in the network. Next, one node *i* is iteratively re-assigned to each of its neighbors’ clusters and the impact on the network’s modularity is assessed. Node *i* is then re-assigned to the cluster where its inclusion results in the greatest gain in modularity. This process is repeated until no gain in modularity can be achieved, that is, a local maximum has been found. In the second step, a new network is constructed from the identified clusters. Edge weights between the nodes, including self-loops, are computed by summing over the weights of the links that connect nodes in each cluster. The first step is then reapplied on the resulting network. These two steps are then repeated iteratively to improve the cluster assignments.

Our aim is to predict regulatory relationships between kinases and as a result our network is directed, that is up to two directed edges connect each pair of kinases, one for each direction of regulation. As this method only clusters networks with at most one edge connecting each node pair, we retained the higher-probability edge of the two linking each pair of nodes. Prior to clustering, we removed regulatory relationships with posterior probabilities less than 0.5 in order to only retain high confidence predictions. The remaining probabilities were then max-min scaled to derive edge scores on the scale 0.0 to 1.0.

In order to determine if the derived clusters reflected known physiological relationships, we tested the clusters for enrichment in pathway annotations from the Reactome database (Fabregat et al., 2018). For the clusters with 10 or more kinases, we tested the relative frequency of pathway annotations of the kinases assigned to the cluster relative to the frequency of those annotations for the entire set of 504 kinases using the hypergeometric test as implemented by the ReactomePA package for R (Yu and He, 2016). We adjusted test *p*-values using the Benjamini-Hochberg method for controlling the false-discovery rate (Benjamini and Hochberg, 1995) and we set a critical value of 0.05 for testing significance. 311 kinases were annotated in Reactome V. 62 (accessed from: http://reactomecurator.oicr.on.ca/download/archive/62.tgz, 13 March 2019) with 6771 pathway annotations altogether.

### Comparison of distances between pathway annotations within and across clusters

We extracted the human protein-protein interaction network from IntAct (version: Oct. 2018) (Orchard et al., 2014). Additionally, on this network, we integrated the human phosphorylation events extracted from SIGNOR, PhosphoSitePlus and OmniPath (Perfetto et al., 2015; Hornbeck et al., 2015; Türei, Korcsmáros, and Saez-Rodriguez, 2016), resulting in a network containing 17089 nodes and 166757 edges. Given a pair of pathway annotations, we computed the mean of all shortest path distances between the proteins annotated for the pair.

These distances were divided into two sets: distances between pathways that are enriched in the same cluster (*n* = 811) and distances between enriched pathways across clusters (*n* = 1019). Furthermore, we excluded distances between pathways that shared kinases, which reduced our within-cluster set to 67. We used the Wilcoxon rank sum test to determine if there was a significant difference in distance between the two sets.

### Comparison of Modularity between Our and Random Networks

To assess the modularity of our network we compared it to a set of randomly generated networks. Our reference network was generated by discarding all edges with probability lower than 0.5. The remaining edges were then min-max scaled to get an edge weight distribution of values between 0 and 1. A set of randomized networks (*n* = 1000) with the same degree distribution as the reference network were generated with the “sample_degseq” function in the igraph package. The “vl” method was used for network generation (Viger and Latapy, 2005). At each randomization, the edge weights of the reference network were shuffled and assigned to the randomized network. These were then clustered as described above. The modularity of the of the clustering was calculated with “modularity.igraph” as implemented in igraph (Clauset, Newman, and Moore, 2004; Csárdi and Nepusz, 2006):

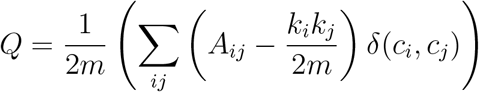

Where *m* is the number of edges in the network, *A* is the adjacency matrix, *k* denotes the degree of the nodes in question and *δ* is an indicator function returning 0 if nodes *i* and *j* are both members of cluster *c*, and 1 otherwise. Applying this procedure to the random networks provided us with an empirical distribution of modularity values from which we derived an empirical *p*-value for the modularity of the reference network.

### Prediction of phosphosite functional sign

As a prerequisite for predicting the sign (activating versus inhibiting) of regulatory relationships, we first built a model to classify individual phosphosites as having either an inhibitory or activating effect on the substrate protein. As features, we used: the percentage position of the site relative to the start and end of the protein kinase domain (i.e. between 0 and 1 for sites that fall within the domain); the percentage position of the site along the protein’s length; the domain (if any) in which the phosphosite lies, including, but not limited to, protein kinase domains; the phosphosite residue (serine/threonine or tyrosine); whether or not the substrate is a tyrosine kinase; an estimate of secondary sequence disorder, as calculated by DISOPRED (Ward et al., 2004); and the −*log*_10_*p*-value of the site being in a phosphorylation hot-spot (Strumillo et al., 2019).

To train and validate our model, we fetched a list of human protein kinase phosphosites annotated as inducing or inhibiting activity from PhosphoSitePlus (Hornbeck et al., 2015). We built our model using BART as described above. We evaluated model performance via 20 iterations of 3-fold cross-validation on a training/validation set of 50 activating and 50 inhibiting phosphosites, sampled randomly each iteration. The final model was trained using the full training set and posterior probabilities of a phosphosite being an activating site were calculated.

We next found the probability cutoff that maximizes the Matthew’s Correlation Coefficient (MCC) for classifying sites as either activating or inhibiting:

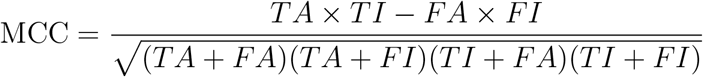

where *TA*, *TI*, *FA*, and *FI* are the numbers of true activating, true inhibiting, false activating and false inhibiting predictions at a given cutoff. Values above the cutoff were taken as “activating” predictions and those below were “inhibitory” predictions. The cutoff that maximizes the MCC was then subtracted from all predicted probabilities, yielding a score of less than zero for “inhibitory” predictions and greater than zero for “activating” predictions. Finally, these scores were rescaled so that the largest absolute value was 1 while maintaining a midpoint at zero.

### Prediction of regulatory sign

We followed a similar procedure for classifying kinase-kinase regulatory relationships as being activating or inhibiting. The predictive features that we used were: the regulator and substrate protein kinase types (serine/threonine versus tyrosine kinases); the signed functional score; a signed formulation of the DCG; and a signed coregulation score. To derive a signed functional score, we simply assigned the sign of the phosphosite sign prediction (negative for “inhibiting”, positive for “activating”) to the site’s functional score. We then used the signed formulation of the substrate’s highest functional score as the final feature.

#### Signed Discounted Cumulative Gain

We modified the DCG calculation to determine whether a regulating kinase tends to “find” inhibitory or activating phosphosites on the substrate kinase. To achieve this, we applied a DCG-like calculation to the signed functional scores, where a positive sum would indicate an activating relationship and a negative sum would indicate an inhibitory one.

If the substrate phosphosites with the highest PSSM scores tend to have high functional scores with the same sign, the initial steps of the DCG cumulative sum will move in one direction. However, if the substrate has many sites and the predicted signs of the sites are unevenly distributed, the sheer number of sites alone would overcome the initial signal from the high PSSM-scoring sites. For example, if the substrate has 3 predicted inhibitory sites which all have high PSSM scores for the regulator and 10 predicted activating sites that have low PSSM scores (but high functional scores), the final DCG on the signed functional scores would ultimately be positive regardless of the site-ordering by PSSM. Thus, we formulated the signed DCG in terms of the most extreme value reached by the sum.

We begin, as with the standard DCG, by ranking the *n* substrate sites according to decreasing PSSM scores (*s*, as described above). This produces an ordered set of their signed functional scores, 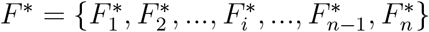. We then calculate a partial DCG on this set, up to the index that produces the largest absolute sum:

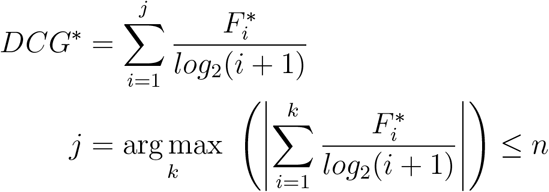

We then normalize *DCG*\* against the most extreme value of the same sign possible for that substrate, retaining the sign. That is, if *DCG*\* < 0 we rank the substrate sites by increasing signed functional score to find 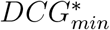, the most extreme negative sum possible for the substrate; and otherwise we rank the sites by decreasing signed functional score to find 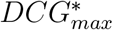, the most extreme positive sum possible. Thus,

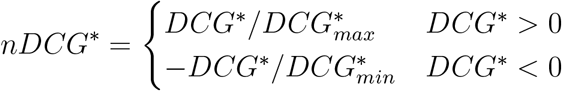

#### Signed coregulation score

In order to produce a signed coregulation score, we followed the same procedure described above for the coregulation score. However, rather than using the *p* value of the correlation test, we used the signed correlation statistic (Spearman’s rho). In order to make the test statistics comparable in spite of differing numbers of data points (i.e. number of conditions or samples in which two phosphosites have both been quantified), we *z*-transformed the scores:

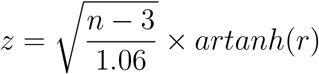

where *n* is the number of coquantified conditions/samples for the pair of sites and *r* is the estimated correlation coefficient. In order to isolate correlations between likely regulatory sites, we then scaled *z* by the signed functional scores of the two sites. Finally, for a given pair of protein kinases, we took the most extreme scaled *z* value as their signed coregulation score.

In calculating these signed coregulation scores, we encountered cases that are inconsistent with a direct regulatory relationship between one protein kinase and another that is governed by a functional phosphosite on the regulator. In particular, in a direct regulatory relationship, the sign of the functional site on the regulator must be the same as the sign of the correlation. For example, if a phosphosite on the regulator is inhibitory (negative), a positive correlation of phosphorylation state with a substrate functional site could only occur through the activity of a third protein kinase (although we note that in kinases with more complicated rules of multi-site regulation, such correlations might be possible). In order to better discriminate strong signals of coregulation, we therefore removed site pairs in which the sign of regulator’s site was incoherent with the sign of the correlation.

#### Training and validation of the sign predictor

We built a predictive model of regulatory sign from these features using BART as described above. As a training and validation set, we gused 503 signed regulatory relationships (394 activating, 109 inhibitory) between protein kinases from the OmniPath database that were supported by at least two data sources. The model was validated via 20 iterations of 3-fold cross-validation, where each iteration used a different random sample of 109 activating relationships for the training/validation set.

We built 20 iterations of the final model using similar random instantiations of the training set. Finally, for each directed kinase-kinase pair, we assigned the mean posterior probability produced by these 20 models as a final regulatory sign score, where a higher value would indicate an activating relationship and a lower score would predict an inhibitory relationship.

## Acknowledgments

B.M.I. received funding from F. Hoffmann-La Roche Ltd.

## Author Contributions

P.B., E.P. and B.M.I. conceived of the study. B.M.I. and B.P. performed the analyses and wrote the manuscript. D.B. and G.G. performed additional analyses.

## Supplemental Tables

- Table S1: Features used to predict phosphosite regulatory signs (Related to Figure 3a)
- Table S2: Phosphosite regulatory sign scores (Related to Figure 3b)
- Table S3: Features used to predict regulatory relationships (Related to Figures 1 and 2)
- Table S4: Kinase-kinase regulatory relationship and regulatory sign predictions (Related to Figures 4 and 5)
- Table S5: Features used to predict kinase-kinase regulatory sign (Related to Figure 3)
- Table S6: Protein kinase cluster assignments (Related to Figure 4e)
- Table S7: Network cluster pathway enrichment (Related to Figure 4e)

## Supplemental Figures

**Figure S1:**
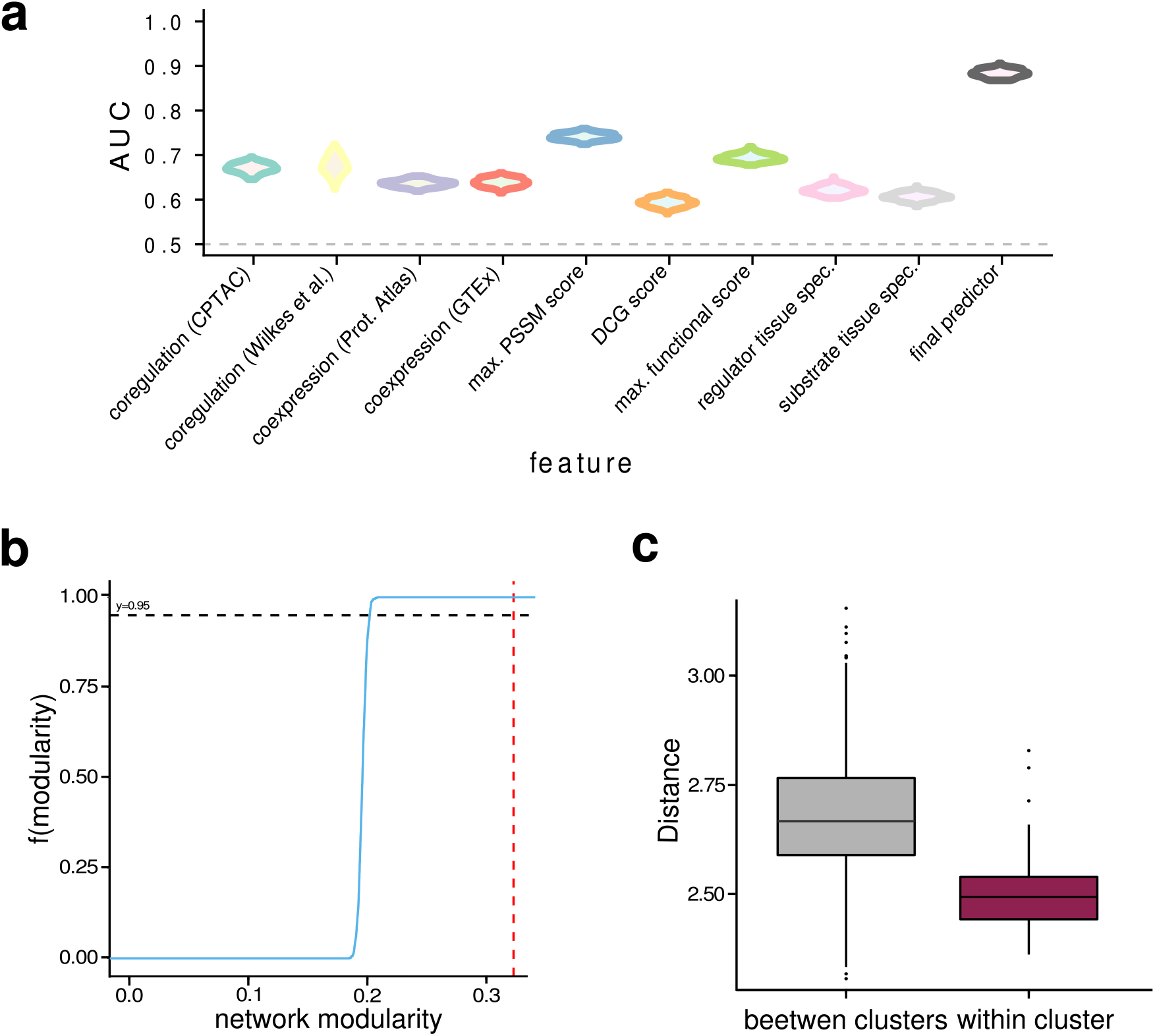
a) Area Under the ROC Curve (AUC) distributions for the edge features and the final predictor across all iterations of cross-validation. b) The modularity of the network (red, dashed line) was significantly higher than expected given random networks with the same degree distribution (cumulative distribution, blue line; the black dashed line shows a cumulative probability of 0.95). c) Pathways associated with the network clusters via statistical enrichment are more closely related to other pathways within the same cluster than with pathways associated with the other clusters.

**Figure S2:**
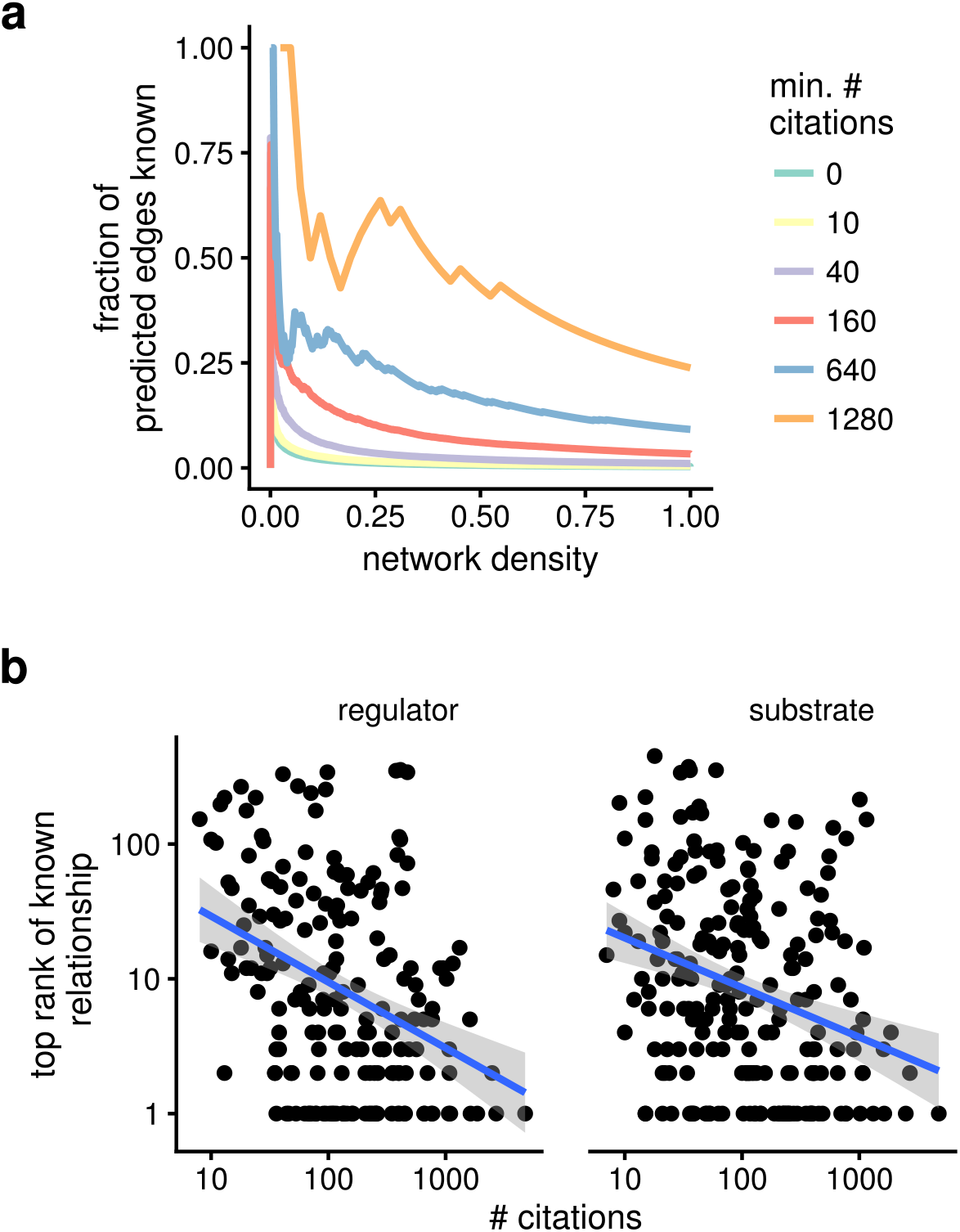
a) Network predictions are dominated by relationships involving under-studied protein kinases. Filtering the network to include only kinases with higher numbers of associated research articles significantly reduces the numbers of novel predictions. b) A correlation between the top rank of a known relationship in the predictions and the number of citations linked to a protein kinase indicates that some study bias is retained.

